# Modeling cellular co-infection and reassortment of bluetongue virus in *Culicoides* midges

**DOI:** 10.1101/2022.06.03.494504

**Authors:** Sean M. Cavany, Carly Barbera, Molly Carpenter, Case Rodgers, Mark Stenglein, Christie Mayo, T. Alex Perkins

## Abstract

When related segmented RNA viruses co-infect a single cell, viral reassortment can occur, potentially leading to new strains with pandemic potential. One virus capable of reassortment is bluetongue virus, which causes substantial health impacts in ruminants and is transmitted via *Culicoides* midges. Because midges can become co-infected by feeding on multiple different host species and remain infected for their entire life-span, there is high potential for reassortment to occur. Once a midge is co-infected, additional barriers must be crossed for a reassortant virus to emerge, such as cellular co-infection and dissemination of reassortant viruses to the salivary glands. We developed three mathematical models of within-midge bluetongue virus dynamics of increasing complexity, allowing us to explore the conditions leading to the emergence of reassortment viruses. In confronting the simplest model with published data, we estimate the average lifespan of a bluetongue virion in the midge midgut is about six hours, a key determinant of establishing a successful infection. Examination of the full model, which permits cellular co-infection and reassortment, shows that small differences in fitness of the two infecting strains may have a large impact on the frequency with which reassortant virions are observed. This is consistent with experimental co-infection studies with BTV strains with different relative fitnesses that did not produce reassortant progeny. Our models also highlight several gaps in existing data which would allow us to elucidate these dynamics in more detail, in particular the times it takes the virus to disseminate to different tissues, and measurements of viral load and reassortant frequency at different temperatures.

## Introduction

Some RNA viruses have segmented genomes, meaning that their genomes are divided into between two (e.g. Lassa virus) and 12 (e.g. Colorado tick fever virus) distinct RNA segments [1]. When two related segmented viruses co-infect a single host cell, virions produced by the cell can consist of segments taken from each of the infecting genotypes, potentially creating a new genotype that is a hybrid of the two infecting strains [1,2]. There are currently 11 families of viruses known to have segmented genomes, including the causative agents of medically important diseases of both humans and animals, such as influenza A virus, Rift Valley fever virus, and bluetongue virus (BTV). Occasionally, reassortment can lead to the emergence of new strains of viruses that more effectively evade host immune responses, often by changing the surface antigens of an already successful strain while leaving other segments intact [1]. These new viruses may then have increased epidemic potential, most famously in the process known as antigenic shift for influenza virus, a major factor in precipitating flu pandemics [3]. Similarly, reassortment can also allow virus strains to more effectively invade new regions or new hosts by generating novel combinations of traits [4].

For segmented viruses that are transmitted by insect vectors, there may be an increased potential for reassortment to occur. Firstly, there is the potential for reassortment to occur in both the primary hosts and in the vector. Secondly, as many vectors feed on different host species [5–7], there is an increased potential for co-infection with strains that typically infect different hosts. Moreover, once infected, insects typically remain infected for the remainder of their lifespan [8], increasing the potential for co-infection during subsequent bloodmeals. Such co-infection can occur either by taking a blood-meal on a co-infected host, or by sequentially taking two or more bloodmeals on hosts infected with different strains.

Despite the high potential for reassortment, there are still within-vector obstacles to a reassortant virus strain emerging – the vector must be co-infected with two different strains, the two strains must co-infect a single cell, viable reassortant virions must be produced by that cell, and these virions must eventually be present in the salivary gland at sufficient concentration for transmission to occur during a blood meal. The vector immune response provides another obstacle to reassortment occurring. Vectors only have an innate immune system, and this is often characterized in terms of ‘barriers’ to infection, including a midgut infection barrier, midgut escape barrier, dissemination barrier, and salivary gland infection barrier [9]. Reassortant viruses or both of their parental strains must successfully navigate each of these barriers and make it to the salivary glands for transmission to occur. Understanding how these barriers interplay with the timing of bloodmeals and other parameters of virus replication is key to understanding how reassortant viruses could emerge in vectors.

In this paper, we seek to understand the interplay between blood meal timing, relative viral fitness, and the length of the incubation period in determining the amount of reassortment that occurs in a vector. We focus on BTV, a ten-segment virus that infects ruminant hosts, in *Culicoides* midges. To this end, we develop a series of three mathematical models of within-midge viral dynamics of increasing complexity, each of which allows us to model a different type of data: infection prevalence, viral load dynamics, and reassortment frequency. The first model describes the midgut infection barrier (MIB), the first barrier to infection a viral infection faces in a vector; the second describes the infection dynamics in a singly-infected midge; and the third describes the dynamics in a co-infected midge, and allows for cellular co-infection and reassortment. By examining the behavior of these models, and comparing them to available data, we highlight the potential effects that a variable incubation period (e.g., due to varying temperature), the length of the gap between infectious bloodmeals, and competition between co-infecting viruses may have on the frequency of reassortment. We also propose new experiments which would generate data that improve our understanding of viral dynamics and reassortment in the midge.

## Methods

### Data

For single-blood-meal infection studies, we largely relied on data from Fu et al. [10]. While there are six studies in total which report viral dynamics following a single BTV infection [10–16], we chose to focus on Fu et al. due to the fact that they reported viral titers, infection prevalence, and dissemination in detail, and that they undertook both oral and intrathoracic infections in parallel. We focused on one study as the viral dynamics are often qualitatively different between studies, for reasons that are unclear (for example, Foster and Jones observed two distinct proliferation phases, while others only observed one [10–12]). Fu et al. infected *Culicoides variipennis* midges with BTV-1, both orally and intrathoracically, and kept them at 24 ± 1 °C. Viral load and infection rate measurements were then made at various intervals up to two weeks. In another arm of the experiment, this study used an additional colony of midges that were refractory to oral infection, though we did not use data from that arm of the experiment.

We used data on reassortment during co-infection from a pair of studies: Samal et al., and el Hussein et al. [17,18] (Fig. S1). These studies were conducted by the same group. Both studies blood-fed *Culicoides variipennis* with BTV serotypes 10 and 17, keeping the midges at 25 °C. Blood meal viral loads in all experiments were ∼6 log_10_ TCID_50_/ml, similar to Fu et al. [10]. They then used electrophoretic analysis of progeny virus from co-infected flies to assess whether isolated plaques were of reassortant viruses or one of the parental strains. The main difference between the studies was that Samal. et al only blood-fed midges simultaneously (i.e. with one co-infected blood meal), whereas el Hussein et al. did this but also looked at sequential infection (i.e. with two separate blood meals separated by a period of time). The periods of time tested by el Hussein were 1, 3, 7, 9, and 11 days following the initial blood meal.

### Midgut infection barrier model

The literature on viral dynamics in insects generally speaks about barriers to infection. These can be either physical barriers, such as the barrier between the midgut cavity and the hemocoel, or immunological barriers, such as when cells utilize RNA interference to inhibit virion production. *Culicoides* midges are typically thought to have a midgut infection barrier (MIB; i.e., a barrier preventing virus particles in the blood meal from infecting midgut cells), a dissemination barrier (DB; i.e., a barrier preventing an infected midge from gaining a fully-disseminated infection), and possibly a midgut escape barrier (MEB; i.e., a barrier to virions escaping the midgut into the hemocoel and the secondary tissues). Other barriers exist in other insects, such as a salivary gland infection barrier and a transovarial infection barrier, but there is no evidence of these in *Culicoides* infected with BTV. Due to the difficulty of distinguishing an MEB from a DB, we model these two barriers as one. It is worth noting that when midges that were refractory to oral infection were instead intrathoracically infected in the Fu et al. study [10], 100% of them were able to transmit through their saliva, compared to 0% when infected orally, highlighting the importance of the midgut infection and escape barriers.

We first focus on the MIB, to gain an understanding of the factors governing this barrier. First, we fitted a logistic function to the prevalence data from Fu et al., to estimate the proportion of midges that have an MIB and the timing of when midges that have such a barrier cease to have detectable levels of virus in their midgut. We fitted a functional relationship,

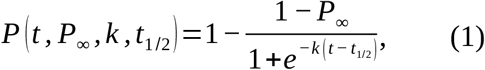

between prevalence, *P*, time since inoculation, *t*, and three parameters, *P*_*∞*_, *t* _1/ 2_, and *k* (Fig. 1). The parameters include *P*_*∞*_, the proportion of midges which achieve a disseminated infection, *t* _1/ 2_, the time when the prevalence is equal to 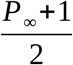, and *k*, which controls the steepness of the curve. Fitting this relationship to data from Fu et al., on the basis of least squares using the optim function in R yielded *P*_*∞*_ =0.346 infected midges per blood-fed midge, *t* _1/ 2_=28.9 hours, and *k* =0.835 hour^-1^.

**Fig. 1:**
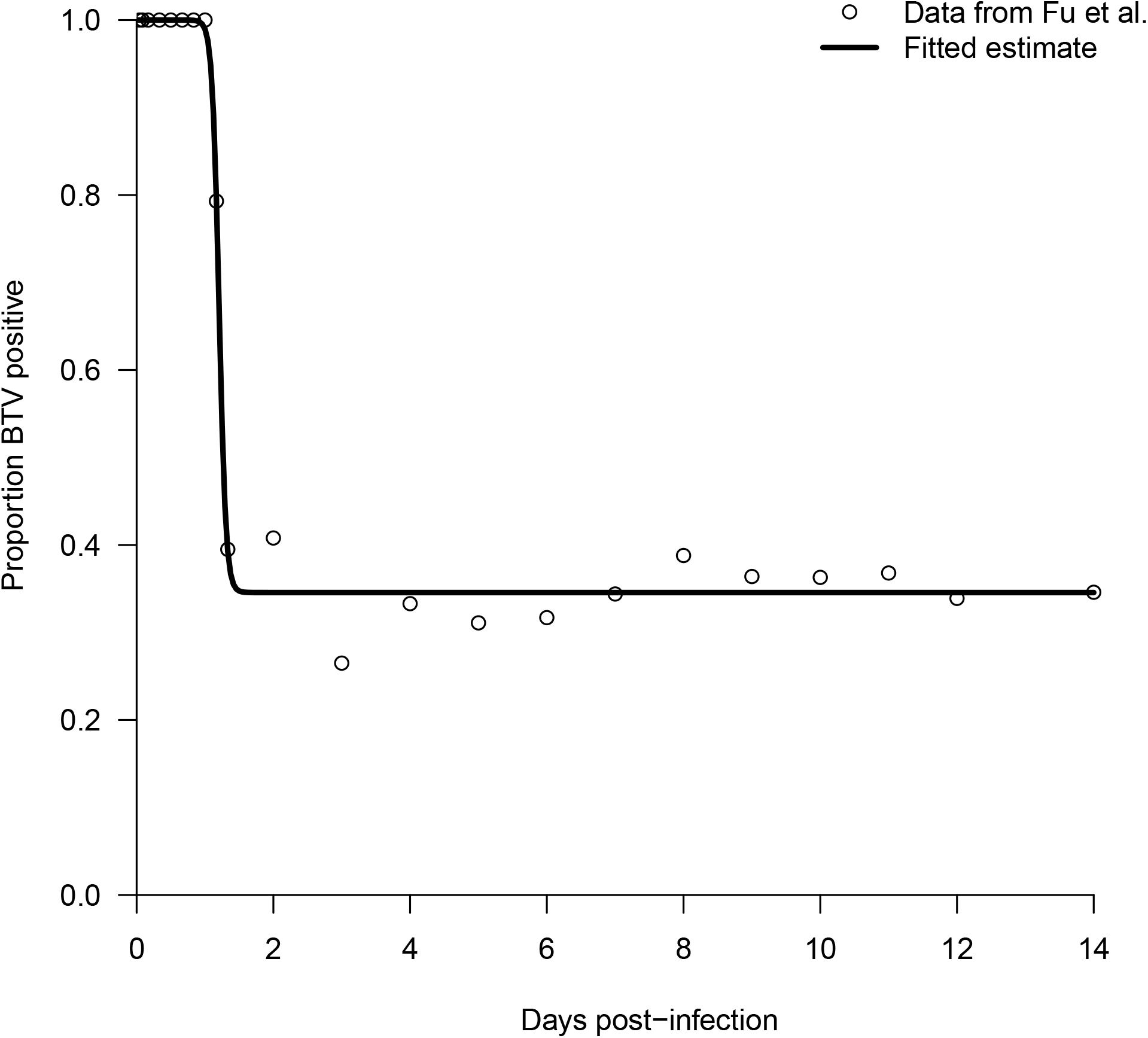
Logistic curve fitted to prevalence data from Fu et al.[10]

Next, we estimated the rate of clearance of virions from the midgut, assuming that they are cleared at a constant rate (Fig. 2). We can model this process as

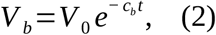

where *c*_*b*_ is the rate virion clearance and *V*_*0*_ is the initial viral load. We then assume that when *t* =*t* _1/ 2_, *V*_*b*_ is at the limit of detection, or *Λ*, because this was the time when half of those midges that did not develop a detectable infection ceased to have detectable virus. Hence, rearranging equation 2, we have

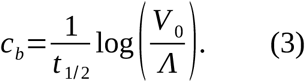

Setting *V* _0_=1260 TCID_50_/midge and *Λ* =10^0.75^, as in Fu et al., implies *c*_*m*_=0.187 virions / hour, or that the average lifespan of a free virion in the midgut is around 5-6 hours.

**Fig. 2:**
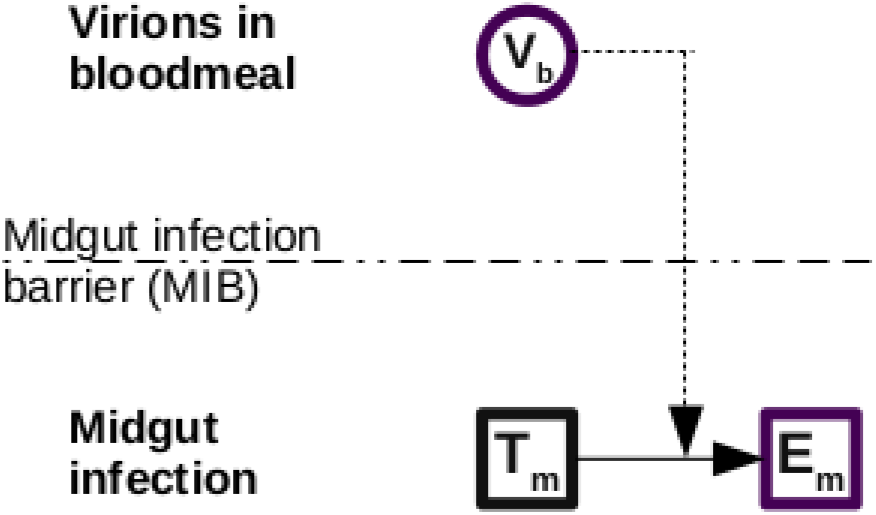
Midgut infection barrier model. The initial blood meal infects susceptible cells in the midgut, if there is no MIB. Virions are cleared at rate c_b_ (not shown). V represents viral load, T uninfected target cells, and E cells that are infected but not yet producing new virus.

There are several different reasons that a viremic blood meal could fail to establish an infection in the midge: a physical barrier to infection; host defenses decrease viral production (e.g., through RNA interference) or increase viral clearance (e.g., via antimicrobial peptides) such that the reproduction number is less than 1; or all viral particles in the blood meal could be cleared before they infect a cell. In the third case, the probability that all virions are cleared before one manages to infect a cell is given by

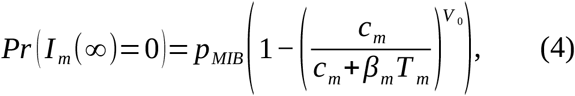

where *p*_*MIB*_ is the probability that the midgut infection barrier is passed given a sufficiently viremic blood meal. This expression allows us to explore the impact of varying blood meal size on the probability of infection, and to distinguish the classical conception of a midgut infection barrier from the stochastic elimination of the viral population.

### Single infection model

We model the infection process and viral dynamics as a system of ordinary differential equations,

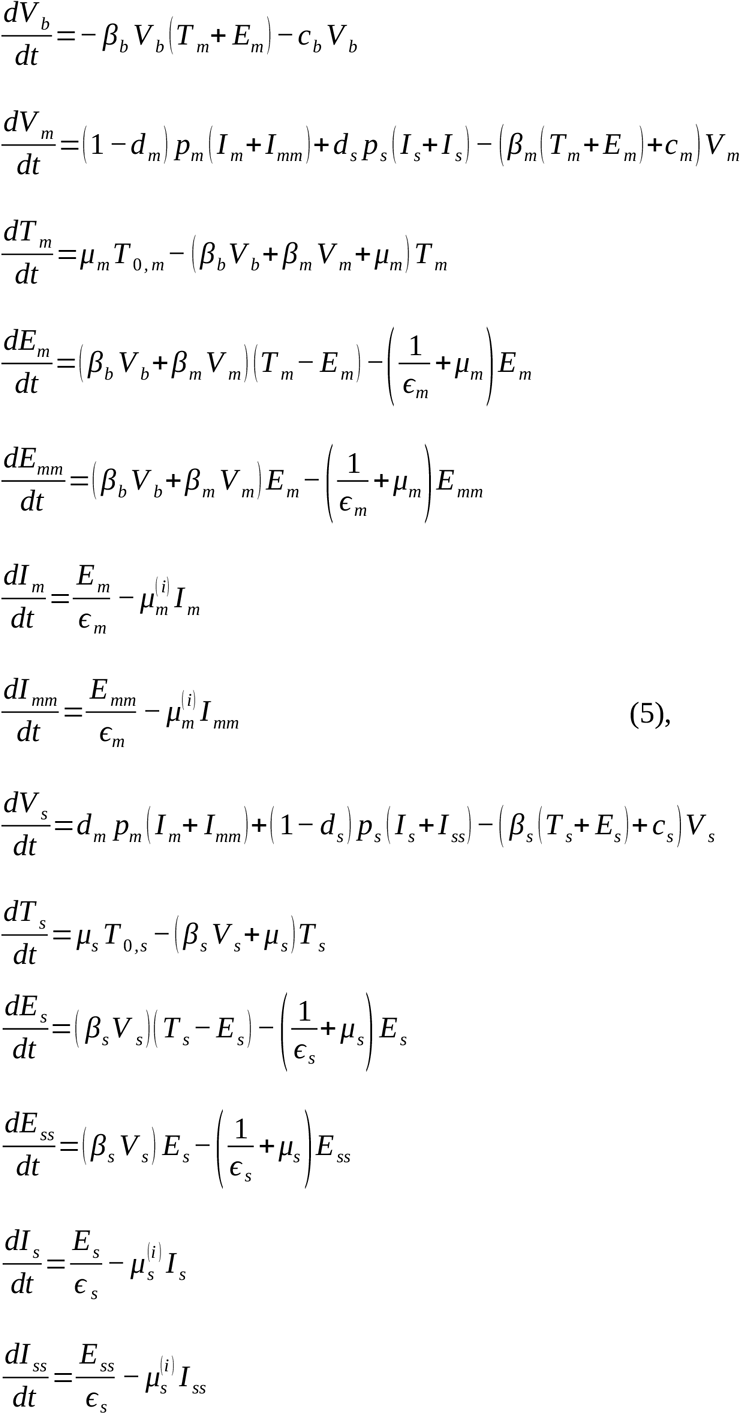

where variables with a *b* subscript pertain to the blood meal, those with an *m* to the midgut, and those with an *s* to the secondary tissues (Fig. 3). State variables with a double subscript (*mm* or *ss*) refer to cells that are co-infected. Allowing for this is necessary to ensure consistency with the co-infection model. In this system, *V*_*b*_ virions are initially ingested in the blood meal. Free virions then infect the uninfected target cells in the midgut (*T*) at rate *β T*. Once infected, a cell enters its eclipse phase, which lasts *ϵ* on average, during which time it does not produce new virions. During the eclipse phase, a cell can become co-infected, in this case only with virions of the same genotype. Following the eclipse phase, cells become productively infected, producing new virions at rate *p*, a fraction *d*_*m*_ of which pass the midgut escape barrier into the secondary tissues. Similarly, for infected secondary tissues, a fraction *d*_*s*_ pass back into the midgut. All cells experience mortality at rate *μ* and virions at rate *c*. New uninfected cells are created at rate *μ T* _0_ to keep a fixed population of cells of *T* _0_. Infected cells may die at a different rate than uninfected cells, though in this study we set these rates to be the same as increased mortality in BTV-infected midges has not been documented.

**Fig. 3:**
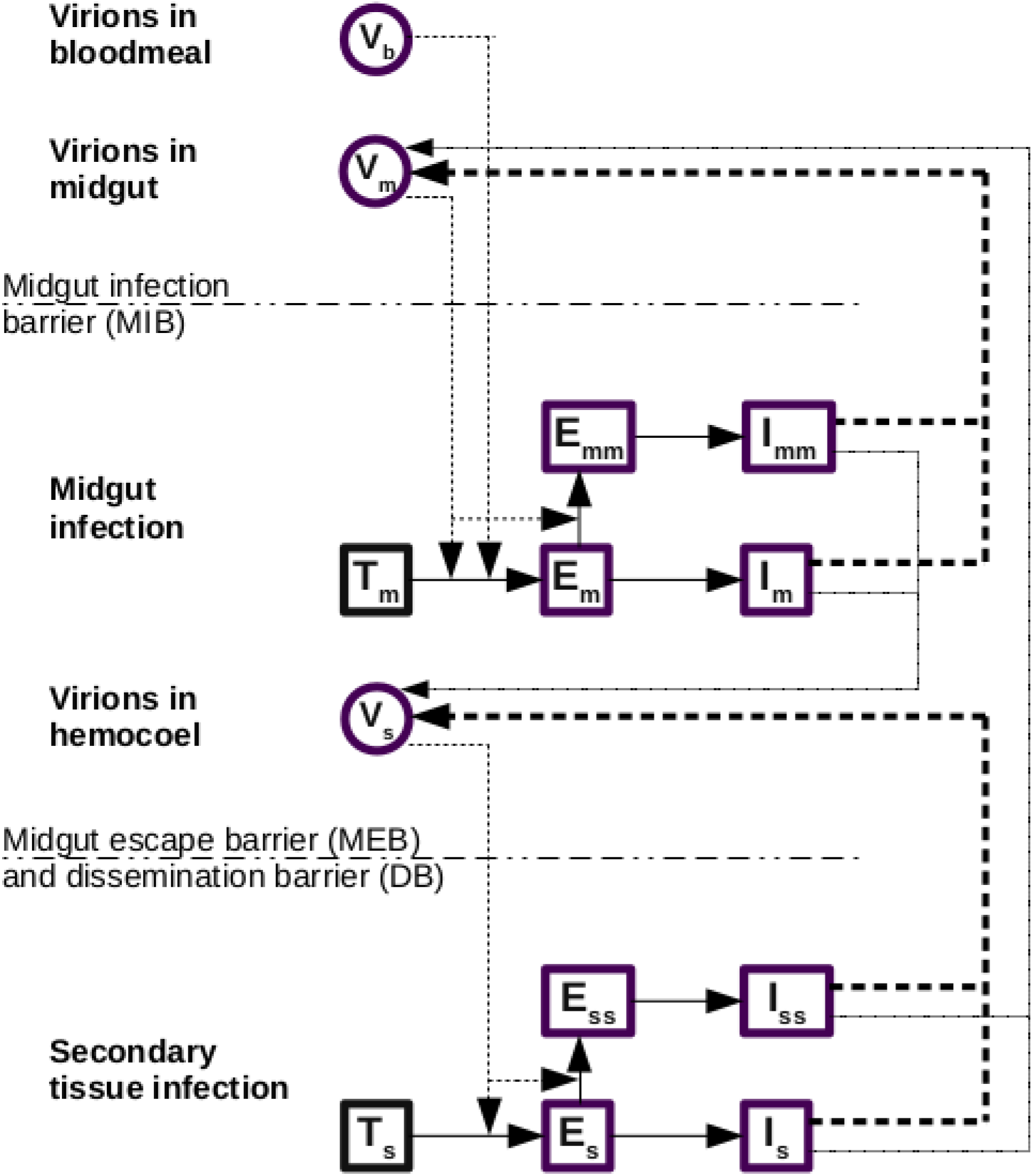
Single infection model. V represents viral load, T uninfected target cells, E cells that are infected but not yet producing new virus, and I cells that are productively infected. Subscripts refer to what region of the body the virus or cells are in (m for midgut and s for secondary tissues), or whether the virus particles were present in the blood meal (b). A double subscript (mm or ss) refers to whether cells are doubly infected; in the single infection model this makes no difference but including these compartments ensures the delay distributions match those of the co-infection model. Solid arrows represent flows, whereas dashed arrows show where rates depend on other state variables. Solid dashed arrows refer to virion production in the same region of the midge, whereas thin dashed line shows produced virions which cross an infection barrier to another region. Infection barriers are shown with a dash-dotted line.

To model the barriers to infection, we simulated the model under three scenarios: 1) with *T* _0, *m*_ =0, to represent midges with an MIB; 2) with *T* _0, *m*_ >0 and *T* _0, *s*_ =0, to represent midges that do not have an MIB but do have a DB; and 3) with both *T* _0, *m*_ >0 and *T* _0, *s*_ > 0, to represent those midges that have neither barrier, resulting in a fully disseminated infection. We then define 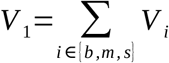 to be the total viral titer of a midge under scenario 1, 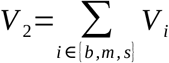 to be the total viral titer of a midge under scenario 2, and 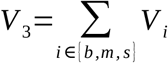 to be the total viral load of a midge under scenario 3. In Fu et al., the average viral load of all positive midges is reported. Initially, this is from all midges, and then once all virions in the blood meal are cleared, it becomes only those midges without a MIB—i.e., those in category 2 or 3 [10]. The average viral load near the start of the experiment, when all midges are infected, will then be

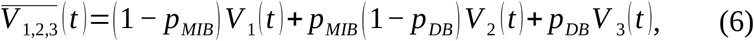

where *p*_*MIB*_ is the proportion of midges that will be infected, and *p*_*DB*_ is the proportion of midges that develop a fully disseminated infection. Similarly, the average viral load near the end of the experiment, when only midges without an MIB are infected, will be:

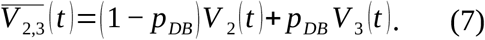

From these, we can calculate the average viral load among positive midges at time *t* as the weighted sum

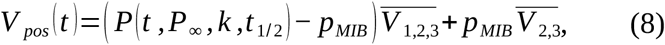

where *P* is the prevalence over time. *V*_*pos*_ is then comparable to the average measured viral load in Fu et al.

### Co-infection model

First, we analyzed how the proportion of reassortant viruses depended on the time since the first blood meal and the gap between the blood meals in the studies of Samal et al. and el Hussein et al. [17,18]. To do this, we fitted a generalized additive model (GAM) to the combined data set with a logistic link, with the proportion of plaques that were reassortant viruses as the dependent variables, and two independent variables: the gap between blood meals, and the time since the first blood meal at which a measurement was taken. As the data on the proportion of plaques which were reassortant was quite noisy, using a GAM allowed us to smooth the relationship between blood-meal timing and reassortment and isolate the most important features. We used four and five knots respectively for the gap between blood meals and the time since first blood meal, based on visual inspection of results obtained with varying numbers of knots.

Next, we extended our compartmental model to accommodate co-infection (Fig. 4, see Supplementary Text for equations). We now have three different types of virions, labelled *i, j* for the two infecting strains of BTV and *r* for reassortant viruses. We don’t distinguish between different segment combinations, grouping all reassortants together for simplicity. In this model, cells can become co-infected with one or two different types of virion, but not three. The second infection can occur at any point during the eclipse phase of a cell, but we assume that once a cell is productively infected, it becomes refractory to further infection. Note that cells infected with a particular type of virion can also be re-infected with the same strain during their eclipse phase, but that cells co-infected with the same virion have no difference in virion production than cells singly infected with that virus. Allowing re-infection in this way ensures that the model achieves ecological neutrality in the sense described by Lipsitch et al. [19]; i.e., the dynamics of the ecological variables (the total number of cells infected with one type, the total number infected with two types, etc.) only depends on the ecological variables, not the specific strains involved. The model also achieves population genetic neutrality, in the sense that there is no *a priori* stable strain equilibrium of strain frequencies and that it instead depends on the initial frequencies (Fig. S2). These two types of neutrality help ensure the model does not have a hidden assumption about the equilibrium level of virus types, and hence of the level of re-assortment.

**Fig. 4:**
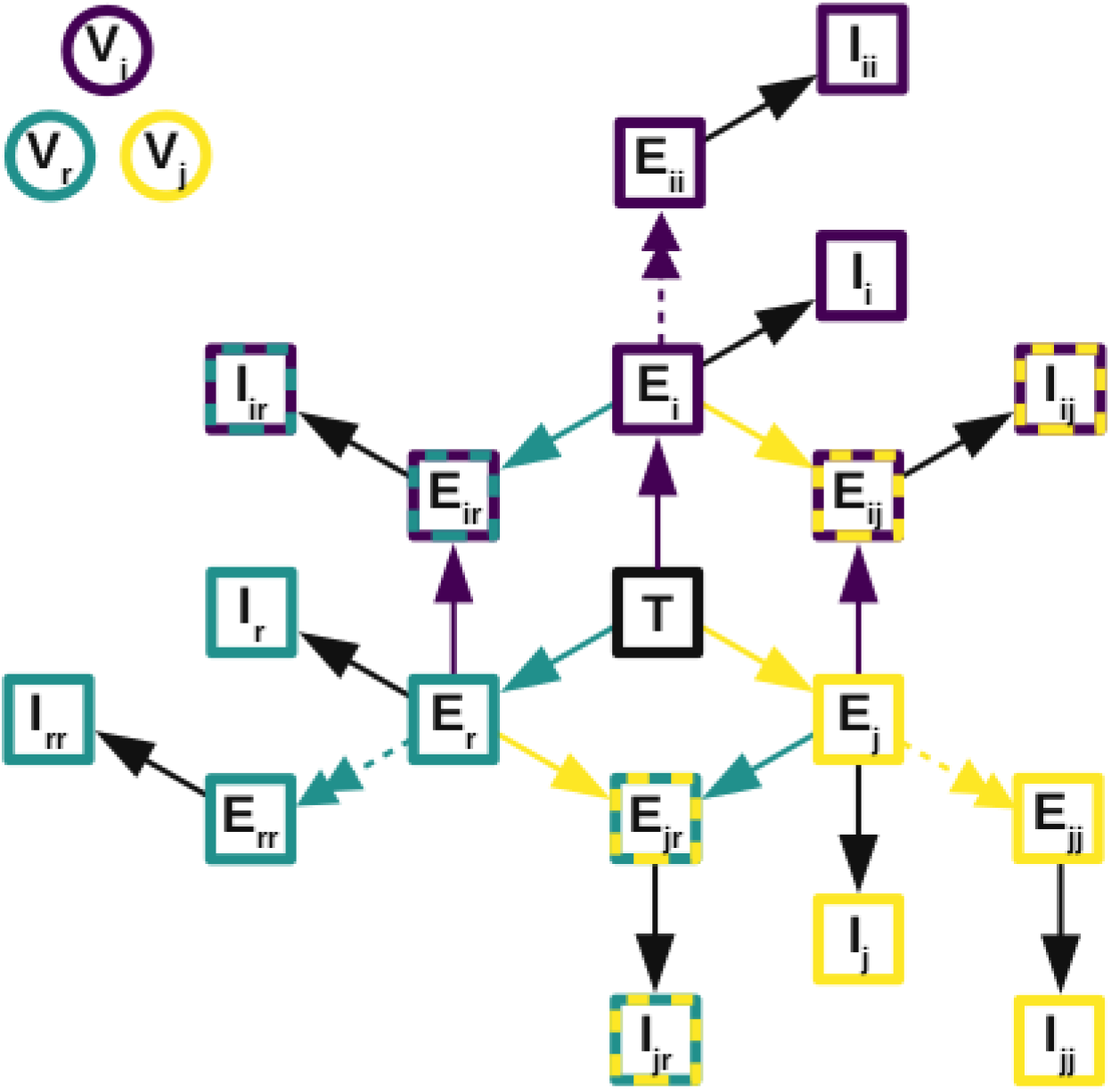
Diagram of co-infection model. Colored boxes refer to the type of virus: purple is virus i, yellow is virus j, and turquoise is reassortant viruses. Where a box is two colors, it indicates cells co-infected with the two indicated viruses. The virus types involved are also indicated by the subscripts inside the boxes. Colored arrows indicate infection with that virus, black arrows indicate progression from the eclipse phase to productive infection, and double-headed dashed arrows indicate when co-infection with the same type of virus occurs. T = target cells; E = eclipse phase cells; I = productively infected cells. Although not shown on the diagram, all cells experience a mortality rate, and new target cells are created, as in the single-infection model.

When a cell is co-infected with two different types of virion, it can produce new virions of either parental type, or reassortant virions. We assume that the proportion of produced virions that are of a particular parental type is

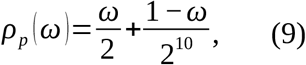

where *ω* is the proportion of produced virions that are with like; i.e., for which reassortment does not occur. This term can be thought of as describing the combined effect of viruses occupying different parts of the cell, for example due to inclusions within the cell [2,20], and incompatible segment combinations. The term 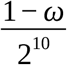 describes those produced virions for which reassortment does occur, but each of the 10 segments is from the same parent. Similarly, the proportion of produced virions that are reassortants is

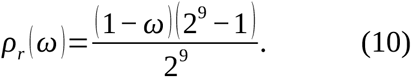

This formulation assumes that all segment combinations are equally likely, and that there is no preference in production for either of the co-infecting virus types. We also assume that the overall rate of viral production is unaffected by the multiplicity of infection. In the absence of reassortment (i.e., when *ω*=1), the co-infection model shows identical viral load dynamics as the single infection model, irrespective of the initial mix of virus types, provided that both virus types have the same parameters (Fig. S2).

The results of Samal et al. and el Hussein et al. [17,18] (Fig. 5) suggest that when there is a small gap between infectious blood meals, more reassortment is observed. We hypothesize that this may be due to a decreased midgut escape barrier when multiple blood meals are taken, consistent with the work of Armstrong et al and Brackney et al [21,22] that suggests that micro-perforations in the basal lamina caused by multiple blood meals aid viral dissemination. We model this as an increase in the probability of passing the midgut escape / dissemination barrier on the logistic scale, to ensure the probability remains less than 1, resulting in

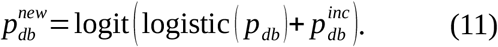

Hence, if we assume that the parameters governing the infection dynamics do not differ between the two infecting viruses and reassortant viruses, then the co-infection model has only two additional parameters compared to the single-infection model: *ω* and 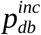.

**Fig. 5:**
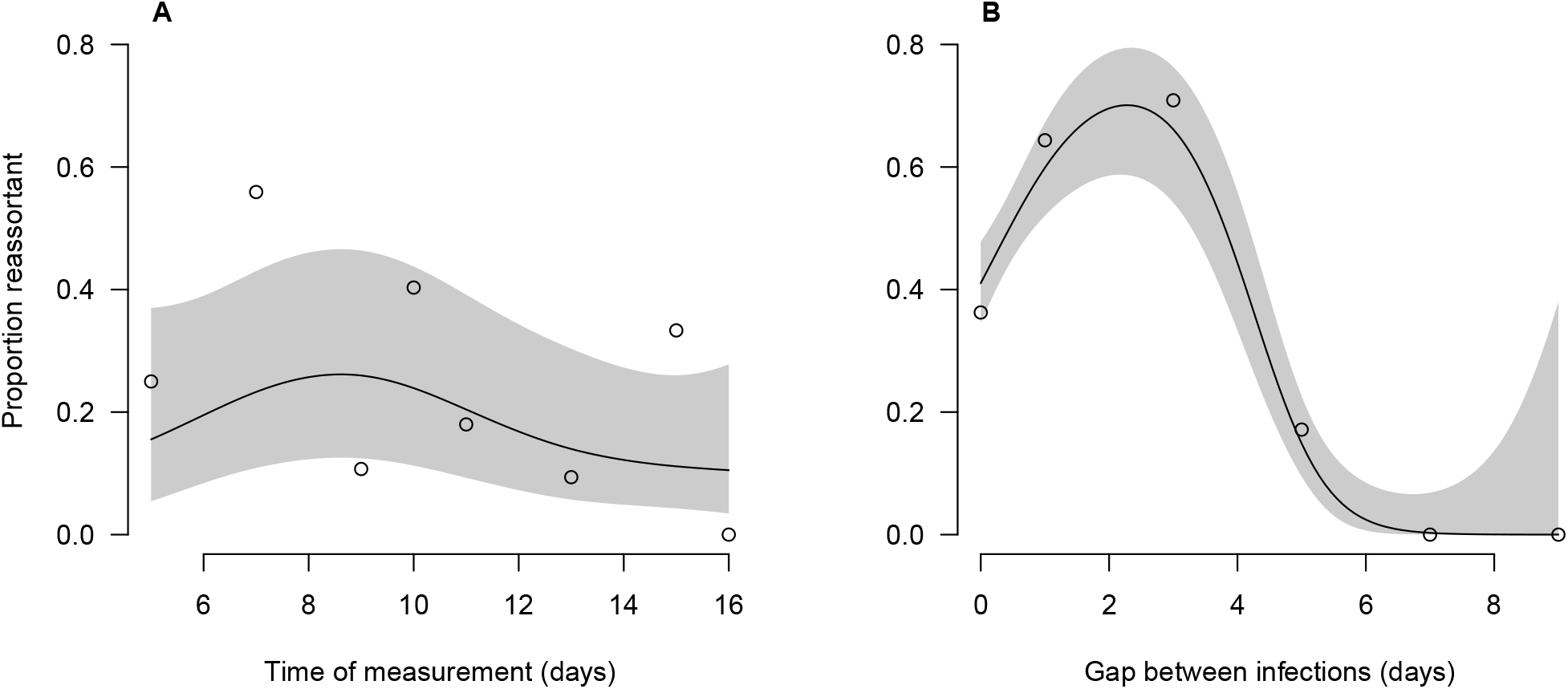
Relationship between the time at which the measurement is taken (A) and the gap between blood meals (B), and the proportion of virions that are reassortant. The line shows a GAM fit to these data, and the shaded region shows the 95% confidence interval.

### Parameter estimation

We jointly fitted the single-infection model and the co-infection model to data using Markov chain Monte Carlo methods. For the single-infection model, we fitted to data from Fu et al. on both oral infection and intrathoracic infection, using a Poisson likelihood,

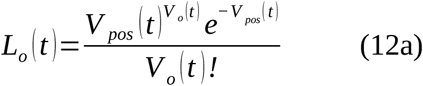

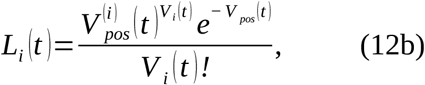

where *V* _*i*_ (*t*) and *V* _*o*_ (*t*) are the viral loads at time *t*, following intrathoracic and oral infection respectively, found by Fu et al., and *V*_*pos*_ (*t*) and 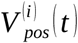 are the total viral loads for a positive midge predicted by the corresponding oral or intrathoracic infection model. We simultaneously fitted the co-infection model with reassortment to the prediction of the GAM based on data from Samal et al. and el Hussein et al. (Fig. 5), using a binomial likelihood,

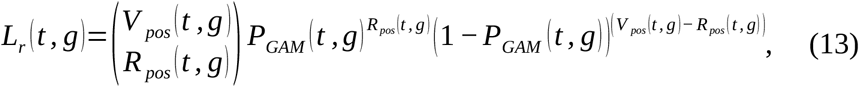

where *V* _*pos*_ (*t, g*) and *R*_*pos*_ (*t, g*) represent the total viral loads and reassortant viral load, respectively, and *P*_*GAM*_ (*t, g*) represents the predicted proportion of virions that are reassortant, at time *t* when blood meals were separated by *g*. We fitted to the GAM prediction as the data are somewhat noisy and we felt this approach may help the dynamical model match the important features of the relationship between blood meal timing and reassortment, particularly the peak in reassortment when the gap between blood meals is around three days. We then summed the log-likelihoods to obtain the overall log-likelihood,

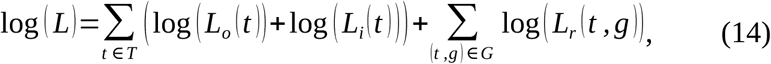

where *T* is the set of time-points used by Fu et al., and *G* is the set of combinations of time-points and gaps between blood meals used in Samal et al. and el Hussein et al. As there is little prior knowledge of the values of any parameters, we used uniform priors on all parameters, with ranges given in Table S1. We used the DEzs MCMC sampler as implemented in the BayesianTools 0.1.7 package in R 4.0.0 [23,24].

### The roles of temperature and viral fitness

Wittman et al. showed that temperature has a large impact on the extrinsic incubation period (EIP) of BTV in *Culicoides*, finding that 1/EIP is linearly related to temperature [14]. The same study also showed that temperature had a negligible effect on vector competence. Additionally, there is evidence that the viral dynamics differ between serotypes, and between strains within a serotype [11,25]. Both of these forms of heterogeneity are likely to have important impacts on the proportion of virions that are reassortants, so we explored their effects on reassortment. Both EIP and the overall dynamics are a complex combination of all parameters, so we made the following simplifications,

a. To assess the role of EIP, we varied the length of the eclipse phase in both the midgut and secondary tissues. This changes the EIP, without affecting equilibrium viral loads, in line with Wittmann et al., who defined the EIP as the first time following the eclipse phase that viral loads reach 10^2.5^ TCID_50_/midge.
b. To assess the role of differing dynamics between serotypes, we varied the ratio of the viral production rates, *p*, between the two viral types involved in the co-infection, while keeping the geometric mean of these parameters fixed at its fitted value. We kept the production rate of reassortant viruses at the baseline fitted level (so it will be between the two production rates of the parental types), but tried an additional sensitivity analysis where it was set to the maximum of the two parental types. Increasing *p* both increases the equilibrium viral load of that virus type and increases its growth rate.

We assessed three targets in this sensitivity analysis: 1) the proportion of virions that are reassortants on day 10 when midges were co-infected on day 0; 2) the proportion of virions that are reassortants on day 10 when midges where infected with the second virus type three days after the first; and 3) the ratio of target 1 to 2, which gives an insight into whether the proportion of virions that are reassortants increases when there is a short gap between infections, as observed in el Hussein et al. [18]

In addition, we also undertook a systematic exploration of parameter space using a Sobol sweep before calculating Saltelli indices for the same three targets. These indices describe the amount of variation in the output that is explained by a particular input. Which parameters were varied and their ranges are shown in Table S1. We used the sensobol 1.1.0 package in R 4.0.0 for this analysis [24,26].

## Results

### Midgut infection barrier

A midge feeding on an infectious blood meal could fail to become infected either because it has an MIB, or because the infecting virions fail to infect a cell before they are all cleared. At initial viral loads similar to those observed in Fu et al. and a number of other studies (∼10^3^ TCID_50_/midge), we found that the overall probability of infection does not substantially differ from the probability that the midge has a midgut infection barrier, unless the rate of viral entry into cells is very low (Fig. 6A). If the initial viral load is much lower than that observed in Fu et al., then the probability of infection could be lower than that suggested by the MIB alone at more moderate rates of viral entry (*β T*_*m*_ *≤*1 new infected cells per hour per virion) (Fig. 6B). However, at values of the rate of viral entry similar to those that we found when we calibrated the model (*β T*_*m*_ ≃5new infected cells per hour per virion, i.e. it takes 0.2 hours on average for a virion to infect a cell), we found that even very low infecting doses would lead to an infection in the absence of a MIB (Fig. 6B). In other words, there is little dose-response effect on the prevalence of infection.

**Fig. 6:**
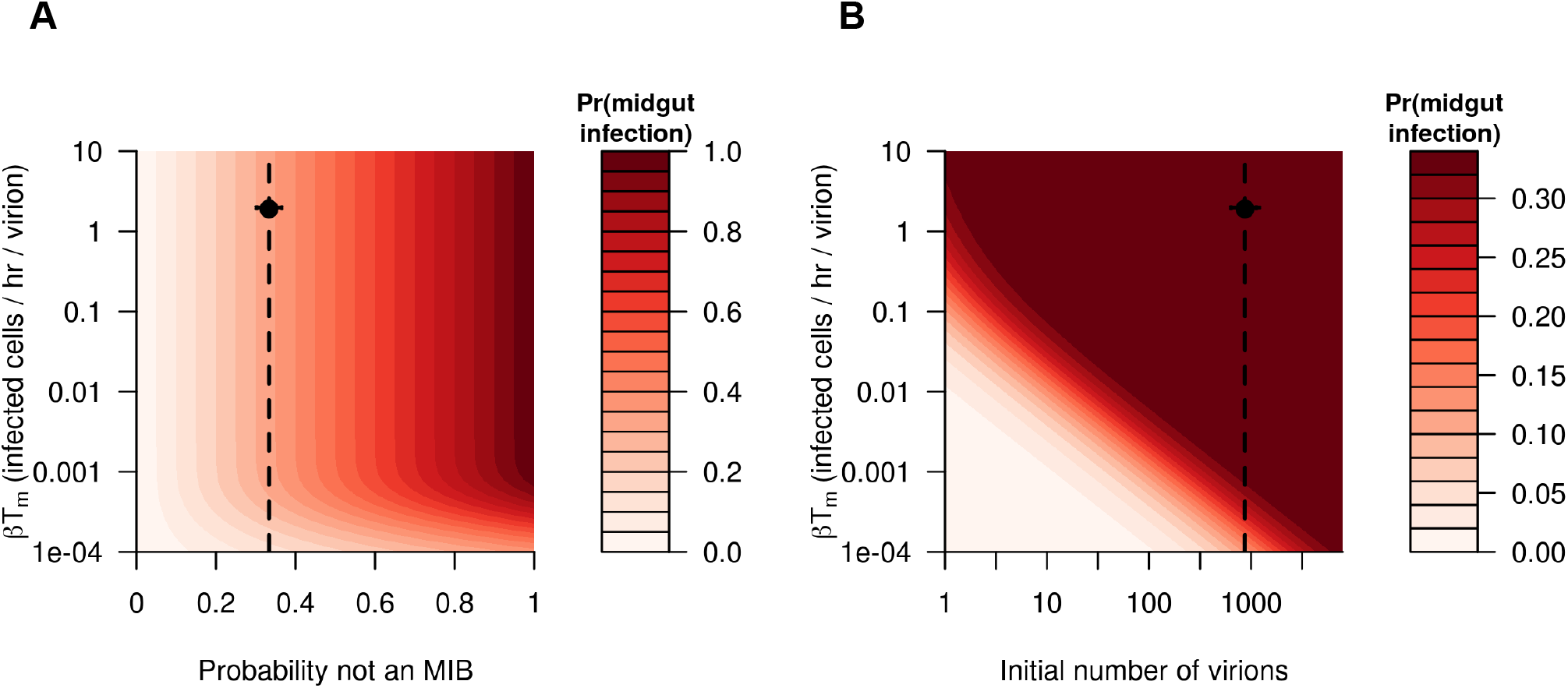
A. The relationship between the proportion of midges which do not have a midgut infection barrier, the rate at which virions enter cells (βT_m_),, and the probability of a midgut infection being established (color-scale), assuming there are 873 virions in the initial blood meal (corresponding to an initial viral load of 1260 TCID_50_/midge). B. The relationship between the initial viral load in the blood meal, the rate at which virions enter cells, and the probability of a midgut infection being established (color-scale), assuming that 2/3 of midges have a midgut infection barrier. In both panels, vertical lines indicate the baseline value of that parameter and circles indicate the best fitting parameters when fitting the model.

### Single-infection model

The single-infection model was able to reproduce several key features of within-midge viral dynamics following both oral and intrathoracic infection. First, the eclipse phase following an infectious blood meal (i.e. the time until the average viral load of positive midges has reached a level higher than the infecting dose) lasted on average about 100 hours, and the viral load reached its minimum value after around 30 hours (Fig. 7A). Our model highlights two differing mechanisms causing this eclipse phase: 1) when a blood meal is imbibed, it takes time before cells are infected and until those cells produce new virions (see black lines in Fig. 7); and 2) early in the experiment, the average viral load of positive midges consists of both infected midges and midges that are not infected but still have virus remaining from the blood meal, so averaging these decreases the measured viral load (compare the black, red, and green lines in Fig. 7A). Second, our model is able to capture the observation of Fu et al. that some midges become infected but do not develop a fully-disseminated infection, thereby remaining positive for the duration of the experiment but with a steadily declining viral load (gray line in Fig. 7A). Our model is also able to reproduce the viral dynamics in those midges which do not have any barriers to infection, although in this case the equilibrium viral load is slightly underestimated (Fig. 7B). This suggests that this population of midges may be distinguished by more than just an absence of midgut and dissemination barriers; for instance they may also have higher rates of viral production in their secondary tissues.

**Fig. 7:**
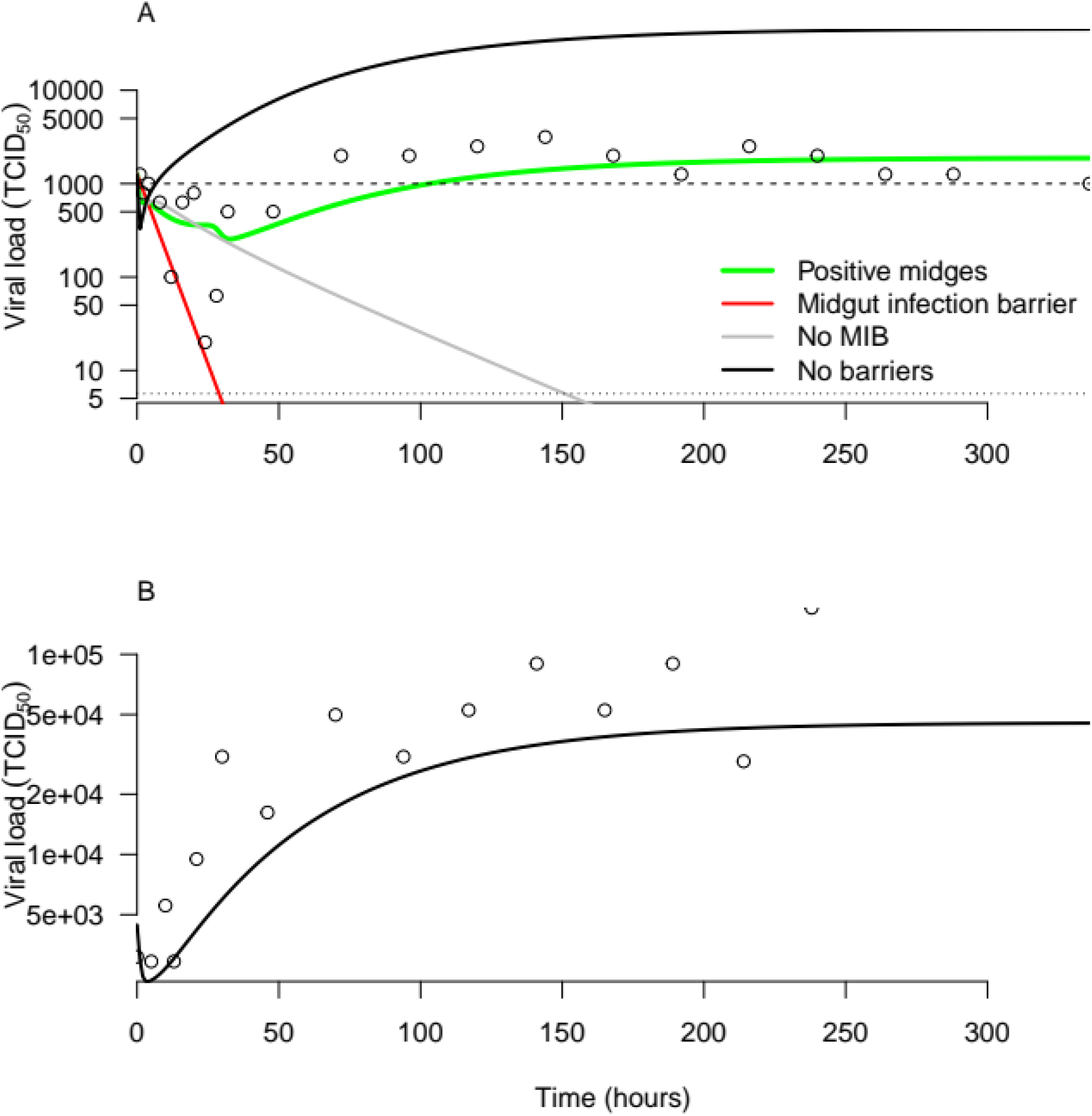
Viral load dynamics following oral infection (A) and intrathoracic infection (B). In both panels, circles show data from Fu et al. A. For the oral infection, the red line represents midges that have a midgut infection barrier and so do not become infected (observed viral load is simply due to virus present in the initial bloodmeal); the gray line represents midges that have no midgut infection barrier but have a dissemination or escape barrier, and so do not develop a fully-disseminated infection; the black line represents midges that have no barriers and so develop a fully-disseminated infection; and the green line is the average viral load of midges that are still positive. The dashed horizontal line shows the initial viral load (1,260 TCID_50_/midge), while the dotted horizontal line shows the limit of detection (10^0.75^ TCID_50_ /midge). B. In an intrathoracic infection, 100% of midges develop a fully-disseminated infection (black line).

### Co-infection model

We jointly calibrated the model to single-infection dynamics and the proportion of virions that are reassortant, assuming identical parameters for the two parameters. Following this, we estimated that 17.5% [95% CrI: 14.4 – 18.2%] of virions produced by a co-infected cell are reassortant viruses, and the remainder are one of the parental virus types. We also estimated that when the midge took two blood meals rather than just one (or in other words, when the two infections were separated in time), the probability that the midge had a DB substantially decreased, from 88% to 2.4% [95% CrI: 2.0% – 2.4%].

The dynamics of total viral load when we modeled co-infection with a single blood meal on day zero, was identical to the dynamics of a single infection (Figs. 7-8, Fig. S3). This is to be expected, as both viruses had the same parameters and the total number of virions per blood meal was fixed, but further verifies that the model behaves in a neutral manner with respect to co-infection. When the second infection occurred later than the first, we observed higher equilibrium loads and an increase in the growth rate around 24 hours after the second blood meal (Fig. 8, particularly for 120-hour gap or more). These effects are caused by the decrease in the probability of a dissemination barrier following the second blood meal, perhaps due to micro-perforations in the midgut basal lamina [21]. The reassortant viral load reached 100 TCID_50_/midge around 24 hours after the second bloodmeal, though this length of time increased as the gap between the blood meal increased (Fig. 8). Notably, it takes much longer (around 48 hours) to reach 100 TCID_50_/midge when midges are fed a single co-infecting blood meal, in part due to the higher dissemination barrier, and in part because there are no pre-existing eclipsing cells as there are when the infections are separated in time.

**Fig. 8:**
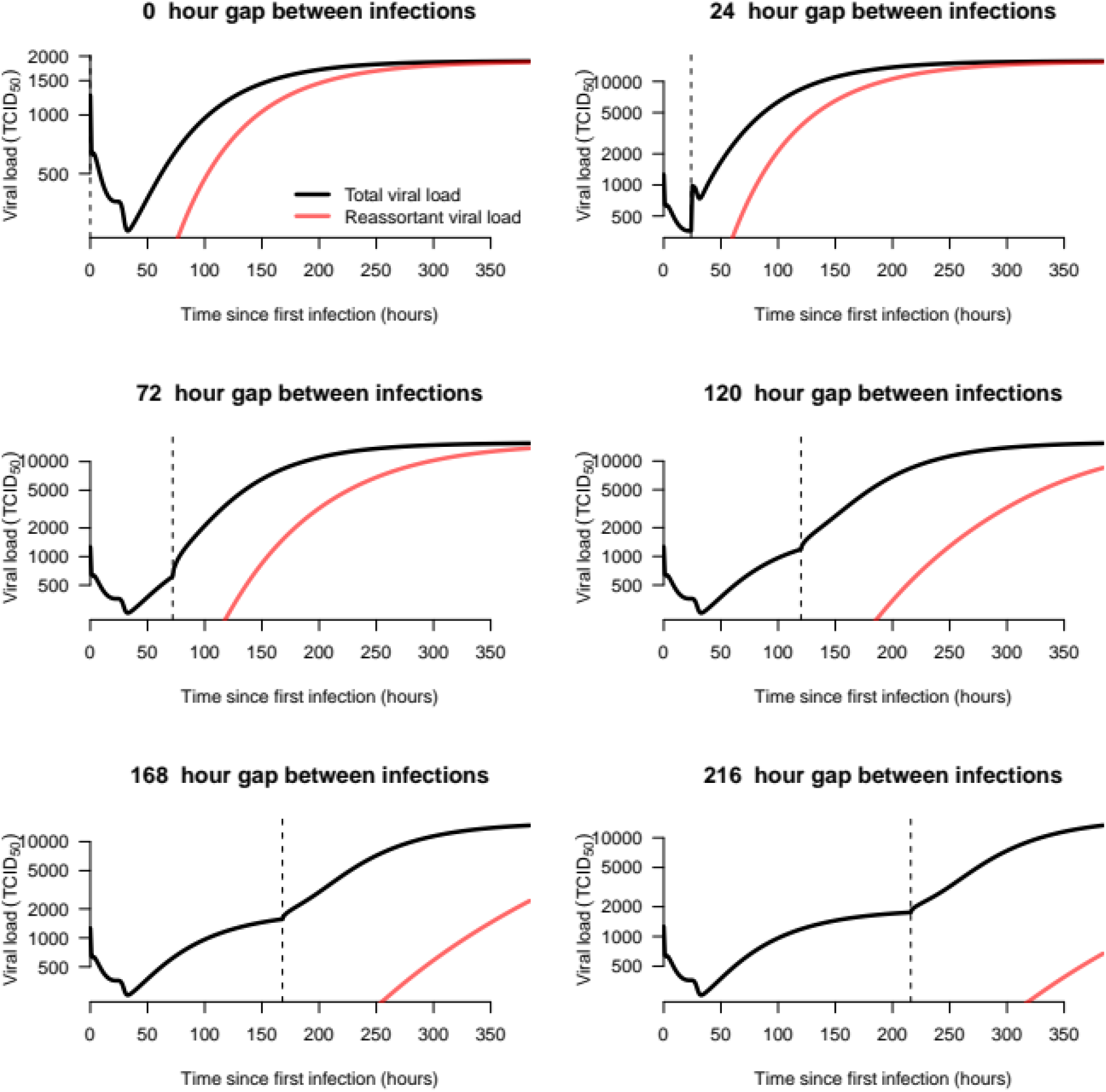
Co-infecting viral dynamics when both viruses have identical parameters. The black line shows the total viral load, and the red line the viral load of reassortant viruses. Each panel represents a different gap between blood meals. The vertical dashed line shows the time of the second blood meal.

In the empirical data, the proportion of virions that were found to be reassortants varied with the gap between blood meals, showing an initial increase to around 70% reassortants as the gap increased from 0 to ∼3 days, before declining to zero at longer gaps (Fig. 5B). There was not a strong relationship between the proportion reassortant and the time at which the measurement was taken, with the GAM fit suggesting the possibility of a slight decrease for later measurements (Fig. 5A). Our dynamical model was able to capture the approximate magnitude of the proportion reassortant for midges that were co-infected with a single bloodmeal on day zero (Fig. 9), and also predicted little or no reassortment for longer gaps. However, the model could not capture the increase in the proportion reassortant as the gap between infections increased to ∼3 days. Although this type of pattern is within the range of the model’s behavior, parameterizations that allowed for this increase were incommensurate with the overall viral load dynamics found by Fu et al. Additionally, our model always predicts increasing proportions of reassortant viruses with later measurements, something which was not found by el Hussein et al. and Samal et al.

**Fig. 9:**
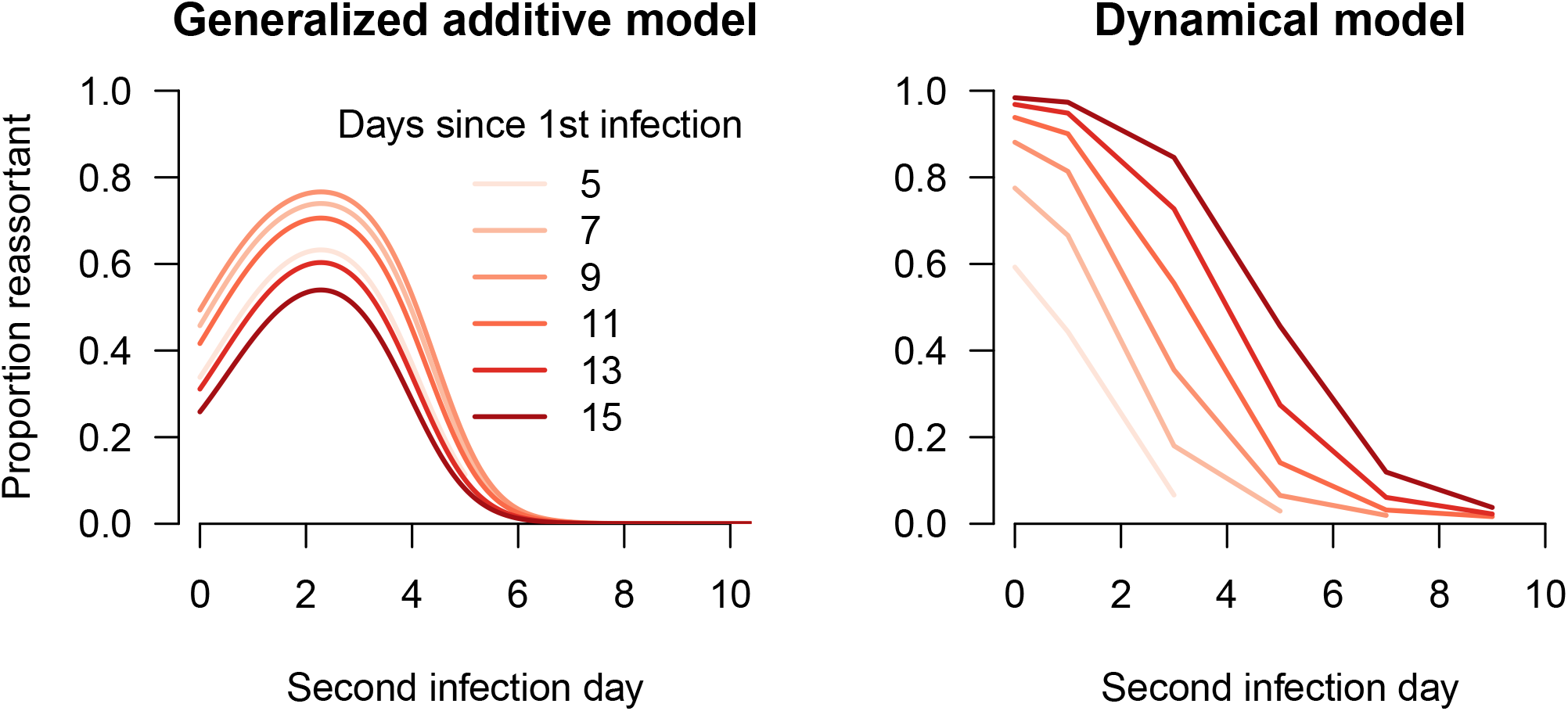
The proportion of virions that are reassortants at different gaps between bloodmeals (x-axis) and different times since the first infection (hue of lines). The left hand plot show predictions from a generalized additive model (GAM) fitted to data from Samal et al. and el Hussein et al, and the right hand the predictions from the dynamical model.

### The role of temperature and viral fitness

To explore the effect of differing viral fitness and a variable incubation period, we varied the eclipse phase and the ratio of the viral production rates between the two viruses, keeping all other parameters fixed. When the two virus types involved with the co-infection had different production rates, we only observed an appreciable amount of reassortment when the production rates were similar (Fig. 10 A, S4). When production rates differed by a factor of two or more, we found that less than 5% of virions were reassortants (Fig. 10 C). This was the case whether it was the first infection or the second infection that had higher production rate. Most reassortment occurred when the second virus had a slightly higher production rate and the eclipse phase was short (Fig. 10 A & C - see the dark red region above the dashed line). When we reduced the eclipse phase, we saw a larger proportion of reassortant viruses by day ten than with the calibrated value of the eclipse phase (Fig. 10 A & B, dotted line). The effect of the eclipse phase on the proportion of reassortment observed depends on the relative viral production rates. For example, if the second virus has a lower production rate, then a longer eclipse phase may lead to more reassortment (Fig. 10, bottom right region).

**Fig. 10:**
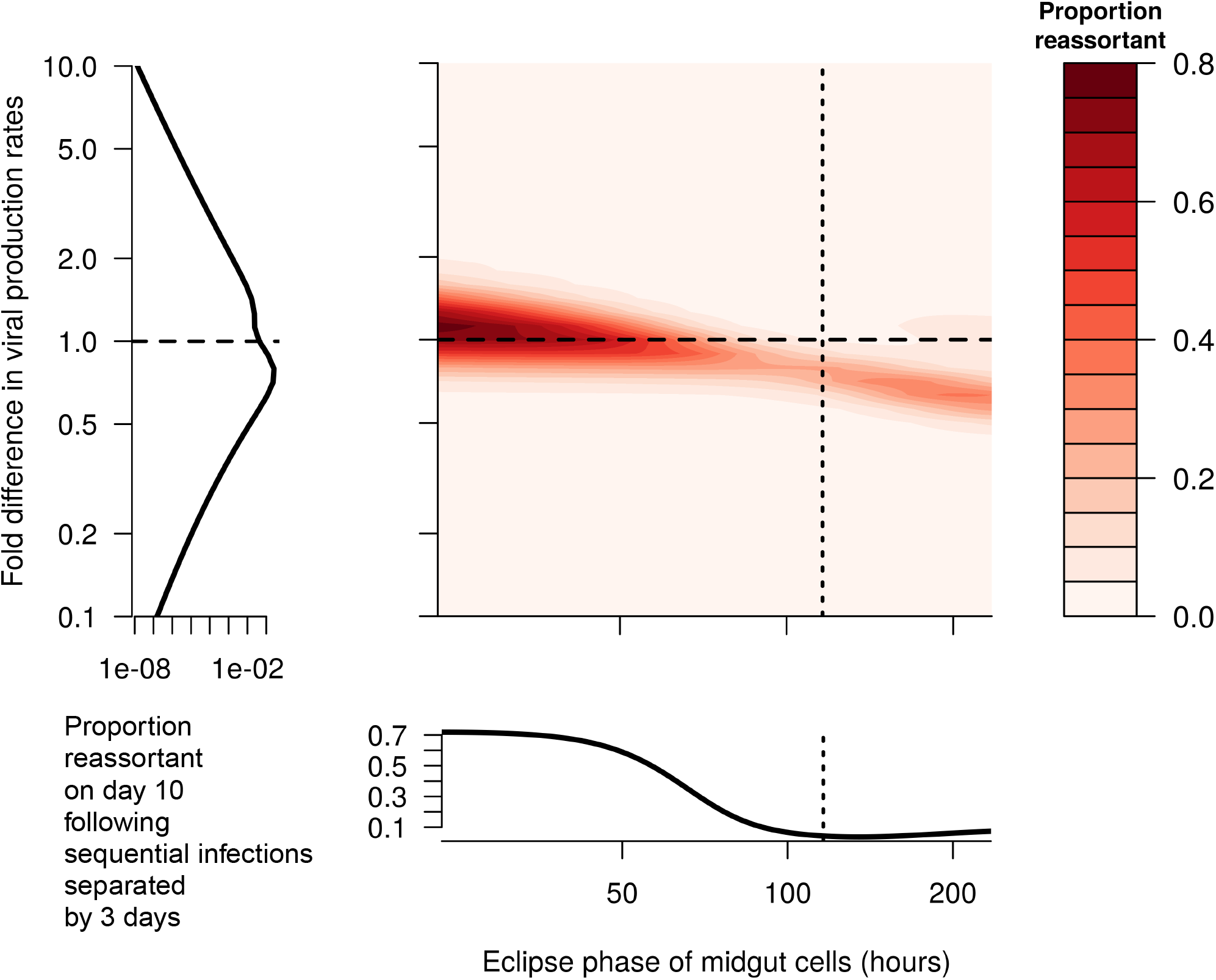
The proportion of reassortant viruses observed ten days after the initial infection (color-scale) when the second bloodmeal occurs three days after the initial infection, for different lengths of eclipse phase (x-axis) and for different viral production rates between the two infections (y-axis). When the fold-change in p is less than 1, the first infecting virus has higher production rate than the second, and when it is greater than 1 the second virus has higher production rate. Dashed lines represent baseline values. Note that the eclipse phase in the secondary tissues was also varied by proportionally the same amount from baseline as the eclipse phase in the midgut (not shown). Note that the left hand plot has an x-axis that is on a log-scale – reassortment decreases very quickly as the difference in viral fitness increases.

When the ratio in the viral production rates was greater than ∼2 (or less than ∼0.5), we were able to recover the increase in the proportion of reassortants observed when the two infections were separated by three days compared to when they both occurred on day zero (Fig. 11, blue region), as indicated by the empirical data (Fig. 5B). However, at most of these parameter values, we also saw very little reassortment in general (Fig. 10), incommensurate with the data from el Hussein et al. The only region of parameter space where there was both substantial reassortment and more reassortment when infections were separated by three days was when the eclipse phase was long and the first virus had slightly higher production rates (Figs. 10 & 11, near the top of the lower right section).

**Fig. 11:**
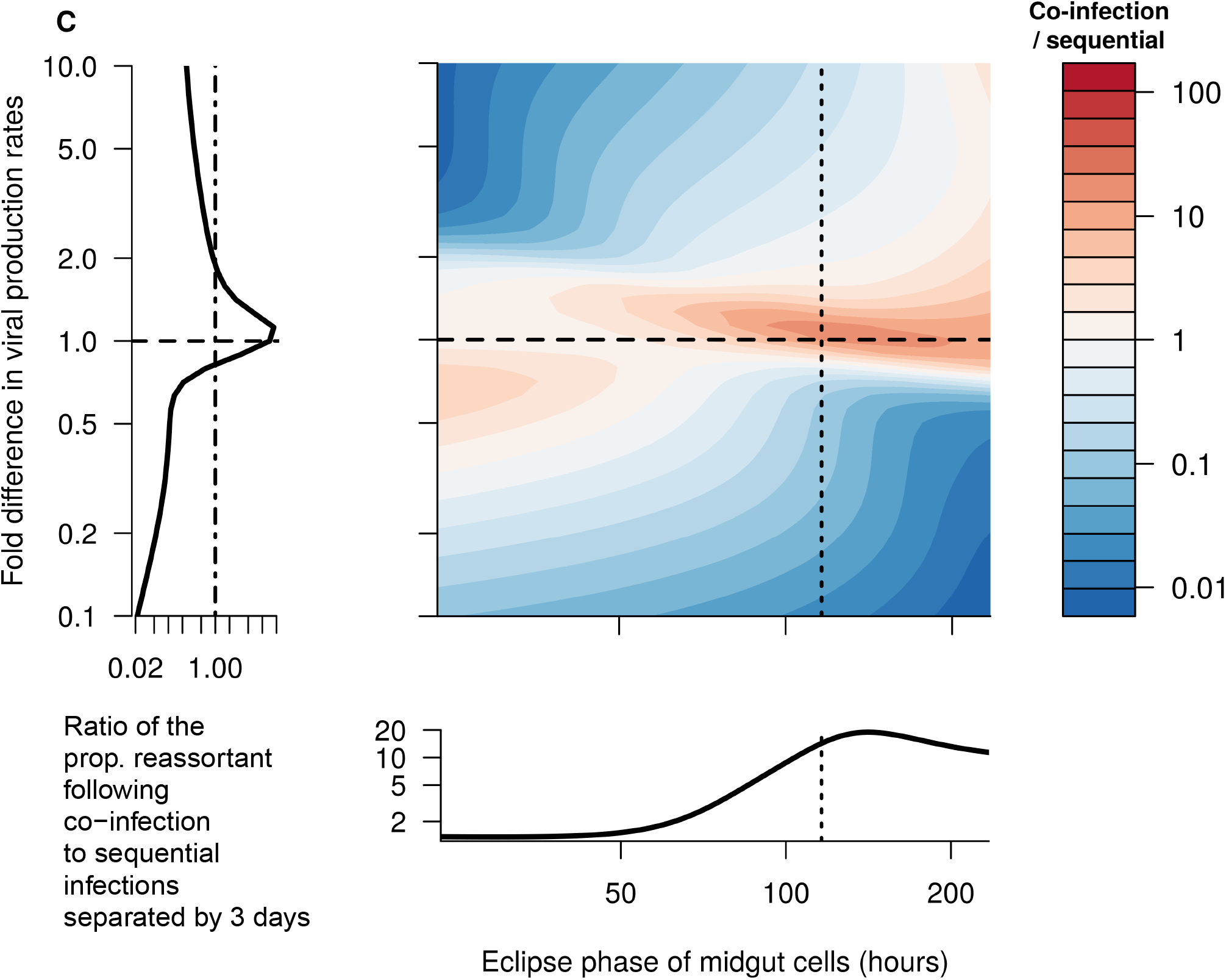
A: The ratio of the proportion of reassortant viruses observed 10 days after the initial infection when both bloodmeals occur on day zero to the same proportion when the second bloodmeal occurs on day three. When this ratio is below 1, it indicates an increase in the proportion of viruses that are reassortant, as observed in el Hussein et al. When the fold-change in p is less than 1, the first infecting virus has higher production rate than the second, and when it is greater than 1 the second virus has higher production rate. Dashed lines represent baseline values. Note that the eclipse phase in the secondary tissues was also varied by proportionally the same amount from baseline as the eclipse phase in the midgut (not shown). B: The ratio of the proportion of reassortant viruses observed 10 days after the initial infection when both bloodmeals occur on day 0 to the same proportion when the second bloodmeal occurs on day 3. When this ratio is below 1, it indicates an increase in the proportion of viruses that are reassortant, as observed in el Hussein et al. Top panel shows changes in the eclipse phase when for equal viral production rates (ie.. horizontal line in A), bottom panel shows changes in the relative production rates holding the eclipse phase fixed (i.e. vertical line in panel A).

In the co-infection model with the calibrated value of the eclipse phase and identical growth parameters for each virus, a variance-based sensitivity analysis suggested that the parameter describing the proportion of produced virions that are with like (*ω*) is most important in determining the proportion of virions that are reassortants overall (Figs. S5-S6). This parameter explains 79% of the variance in the proportion of virions that are reassortants when both infections occur on day zero, and 39% when the second infection occurs on day three. Other important parameters are the rate at which cells die (*μ*) and the rate at which virions are cleared in the secondary tissues (*c*_*s*_). For co-infections, *μ*_*m*_ explains 5.7% of the variance, *μ*_*s*_ explains 1.2%, and *c*_*s*_ explains 0.13%, while for infections separated by three days, *μ*_*m*_ explains 15.4% of the variance, *μ*_*s*_ explains 7.5%, and *c*_*s*_ explains 2.7%. Interactions account for 14% of the variance for co-infections, and 33% for infections separated by 3 days.

## Discussion

In this paper, we used three models of increasing complexity to elucidate the viral dynamics and emergence of reassortment in an insect vector. The first model allowed us to estimate that virions in the midgut have a lifespan of around six hours on average. The second model is able to capture the temporal pattern of viral load within the midge, but is poorly identifiable, suggesting more data is needed to constrain elements of the model. Experiments to obtain such data are proposed below. Our final model captures the broad patterns in the frequency of reassortment at different times since the first infection and gaps between the two infections, but when virus parameters are the same across virus genotypes, it cannot capture the specific pattern of increased reassortant frequency at small gaps between bloodmeals. We were able to recover this pattern when viral production rates of the infecting strains differed, however. The final model also showed that even small relative differences in viral production rates can cause large differences in the frequency of reassortment.

Our results highlight the importance of the relative fitness of the infecting strains and bloodmeal timing. As previously mentioned, we could only reconstruct the increase in reassortment when bloodmeals were separated by three days by allowing the strains to have different production rates. We initially tried to reconstruct this increase by allowing second blood-meals to have a higher probability of passing the MIB, something previously observed in *Aedes aegypti* mosquitoes [22], but surprisingly this could not recreate the observed increase in reassorment when viral production rates were identical. Prior studies of BTV co-infection have found conflicting results reagarding reassortment frequency; while both Samal et al. and el Hussein et al. found substantial levels of reassortment when coinfecting *culicoides variipennis* with BTV-10 and BTV-17, a more recent study by Kopanke et al. observed no reassortment when infecting *culicoides sonorensis* with BTV-2 and BTV-10 [17,18,25]. A possible explanation for this discrepancy is suggested by our observation that when viral production rates varied by more than a modest amount, very little reassortment was observed. These experimental results and our modeling results suggest that whether reassortment is observed, and how much, may be a consequence of delicate balance between relative viral fitness and the timing of infections.

Our study adds to a growing body of work on within-vector modeling, and in particular contributes a mechanistic description of viral co-infection and reassortment. Previous mechanistic within-vector modeling work has often focused on malaria parasites in *Anopheles* mosquitoes [27–30]. All four studies modeled transitions between the life-stages of malaria parasites in the mosquito, or a subset of them, and did not explicitly model infection of mosquito cells. Prior models of viral dynamics in vectors have focused on Zika virus dynamics in *Aedes* mosquitoes [31,32]. Both of these models modeled virus dynamics phenomenologically as logistic growth, though Tuncer et al. included a semi-mechanistic component describing both midgut and secondary tissues and transport between them. Like the malaria models but unlike our model, neither study directly modeled the infection of mosquito cells. These modeling approaches, though suitable for the questions asked in those studies, are less conducive to modeling cellular co-infection and reassortment, as was our aim here. At least two non-vector models of within-host reassortment have been developed for influenza [33,34]. Koelle et al. proposed a parsimonious compartmental model of cellular co-infection and reassortment. In their model, all co-infected cells produce only reassortant viruses, meaning that eventually all virions are reassortant.

Our study has at least four limitations. First, our model did not account for stochastic effects, which may be important if there is a bottleneck following one or more of the barriers that leads to low viral population sizes. Second, we did not allow for inter-midge variability in viral dynamics, which would allow us to capture the distribution in viral loads and reassortant frequencies. Third, we made the simplifying assumption that reassortants had a production rate equal to the geometric mean of their parental viruses. If all reassortant viruses are much less fit than their parental strains, we would obviously expect much lower reassortant frequency, and this may provide an alternative explanation to the findings of Kopanke et al. [25]. Fourth, we implictly assume that all segment combinations are equally likely, with the exception of the parental combinations whose frequency is determined by the withlike parameter, *ω*. However, as we don’t explicitly track different segment combinations, our reassortment compartment could be thought of as a frequency-weighted sum of all reassortants. The latter two limitations are both partially addressed by *ω*: both reduced reassortant fitness and a reduced set of possible segment combinations could be interpreted as an increase in *ω*.

Our modeling suggests several *in vivo* experiments which could help improve future modeling efforts and in turn our understanding of BTV dynamics in *Culicoides*. First, it would be useful to know the distribution of the timing of dissemination to different body parts, especially the midgut cells, the fat body cells, the ganglia, and the salivary glands, including the time virus is first observed or crosses some threshold in that body part, or the viral load in different tissue types over time. The much smaller size of the midge has made this more difficult for BTV in *Culicoides*, though Fu et al. did record this for a subset of midges without measuring the full distribution of timings [10]. Second, the dose-response effect on viral load, infection prevalence, and reassortant frequency could help inform the model’s structure. For example, if we were to compare the dose-response effect in the prevalence of infection in midges, then we could constrain the rate of cellular entry by virions in the midgut (Fig. 6B). Third, elucidation of the mechanisms governing the barriers to infection (i.e. the vector immune response) would allow a more mechanistic treatment of the barriers. Our model currently treats the barriers phenomenologically, without directly incorporating them into the model structure. Fourth, to account for inter-midge variability and include stochasticity, it would be useful to know the distribution of viral loads across midges over time, and how this differs between those midges which develop a fully-disseminated infection and those which do not. This has been characterised for dengue virus in *Aedes aegypti* [35]. Finally, while we explored one possible effect of different temperatures via examination of a varying eclipse phase, a thorough description of viral load trajectories and reassortant frequencies at different temperatures would enable our more systematic inclusion of temperature dependence in viral dynamics.

Our study adds a mechanistic description of cellular co-infection and reassortment in an insect vector. Our results give quantitative understanding of the typical time a virion is resident in the midgut, and highlight how the amount of BTV reassortment observed during midge co-infection will depend on the interplay between the relative fitness of the two infecting strains and the timing of the respective bloodmeals. The relatively simple approach we employ would be straightforward to adapt to other vector-virus combinations with different barriers to infection, or with barriers of different strengths. It would also be straightforward to incorporate new data types, and we suggest a range of experiments to generate such data. Such experimental work would allow us to better constrain the model’s parameters and relate them to different temperature regimes. In addition, future modeling studies could focus on coupling this model with an epidemiological model to try and understand the epidemic potential of reassortment in the midge, or applying this work to other vector-borne segment viruses.

