## Supplementary table and figures for "Modeling cellular co-infection and reassortment of bluetongue virus in *Culicoides* midges"

**Supplementary tables**

Table S1: Model parameters included in calibration and sensitivity analysis, the ranges explored (i.e. limits of the uniform priors for the calibration), and calibrated values (i.e. *maximum a posteriori* values and 95% credible interval)

| **Symbol** | **Definition** | **Units** | **Explored range** | **Calibrated values (MAP; 95% CI)** |
|---|---|---|---|---|
| $\beta_m T_{0,m}$ | The rate at which virions enter uninfected target cells | $hour^{-1}$ | [5x$10^{-4}$, 5] | 1.90;<br>[1.88, 2.04] |
| $d_m$ | The proportion of virions produced in the midgut which pass the MEB | - | [0, 1] | 0.385;<br>[0.385, 0.495] |
| $p_m$ | The rate at which productively infected midgut cells produce new virions | virions $cell^{-1}$ $hour^{-1}$ | [1, $10^4$] | 194;<br>[194, 225] |
| $T_{0,m}$ | The initial number of target cells in the midgut | cells | [1, $10^4$] | 4010;<br>[3830, 4030] |
| $\mu_m$ | The midgut target cell death rate | $hour^{-1}$ | [0.01, 1] | 1.00;<br>[0.975, 1.00] |
| $\varepsilon_m$ | The average length of the eclipse phase in midgut cells | hours | [0.0167, 120] | 116;<br>[111, 116] |
| $c_s$ | The viral death rate in the secondary tissues | $hour^{-1}$ | [0.01, 1] | 0.448;<br>[0.432, 0.590] |
| $\beta_s$ | The per-virion rate of effective contact between cells and virions in the secondary tissues | $hour^{-1}$ $cell^{-1}$ | [$10^{-7}$, $10^{-5}$] | 8.63x$10^{-7}$;<br>[0.817, 1.40]x$10^{-6}$ |
| $d_s$ | The proportion of virions produced in the secondary tissues which return to the midgut | - | [0.01, 0.99] | 0.138;<br>[0.129,0.141] |
| $p_s$ | The rate at which productively infected secondary tissue cells produce new virions | virions $cell^{-1}$ $hour^{-1}$ | [1, $10^4$] | 9990;<br>[5160, 9980] |
| $T_{0,s}$ | The initial number of target cells in the | cells | [1, $10^4$] | 5360;<br>[4930, 5390] |

| | | | | |
|---|---|---|---|---|
| | secondary tissues | | | |
| $\mu_s$ | The secondary tissue target cell death rate | $hour^{-1}$ | [0.01, 1] | 0.996; [0.751, 0.999] |
| $\varepsilon_s$ | The average length of the eclipse phase in secondary tissue cells | hours | [0.0167, 120] | 96.6; [95.7, 101] |
| $\omega$ | The proportion of reassortants produced by co-infected cells that are withlike | - | [0, 1] | 0.54; [0.504, 0.626] |
| $p_{DB}^{inc}$ | The increase on the logistic scale in the probability of the dissemination barrier being crossed following a second bloodmeal | logistic(virions $cell^{-1}$ $hour^{-1}$) | [0, 10] | 9.38; [9.31, 9.53] |

### Supplementary figures

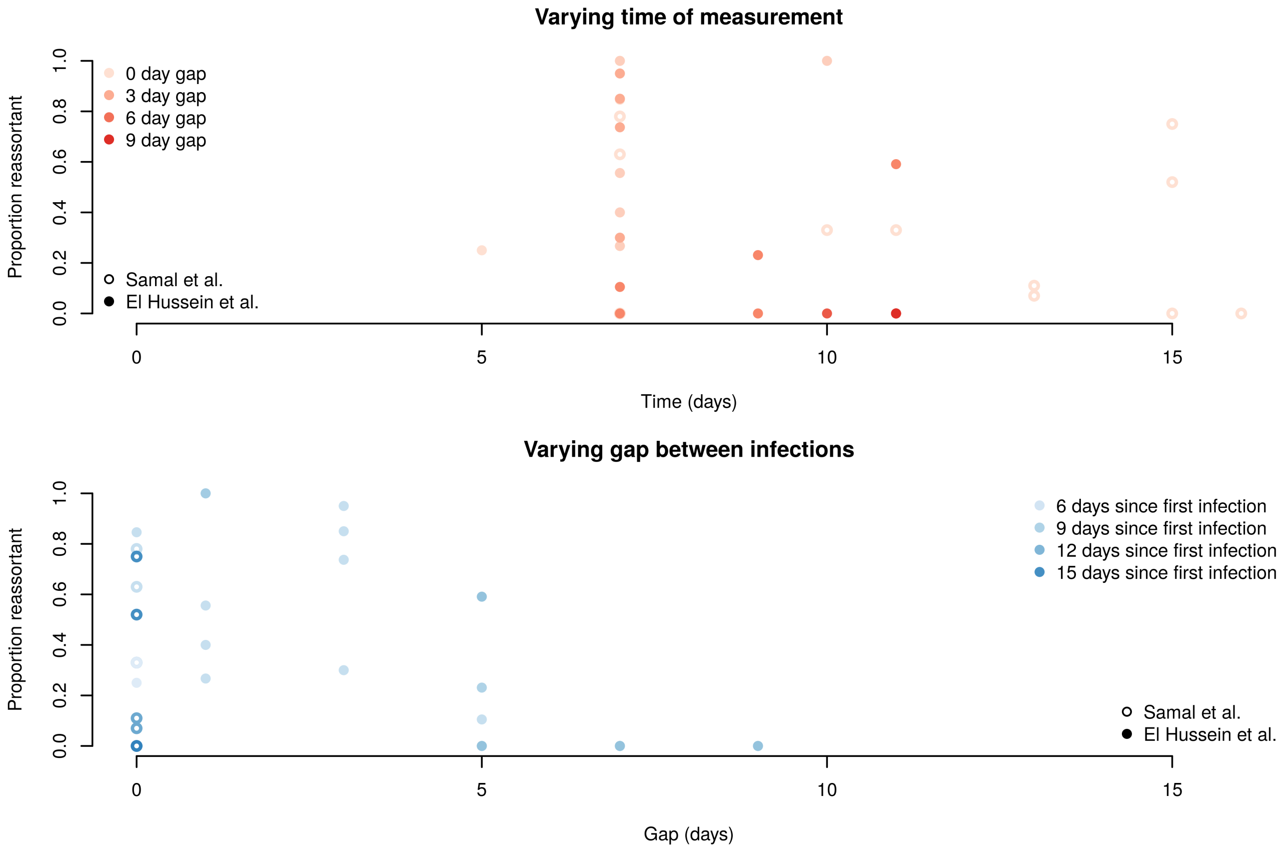


*Fig S1: Data on proportion of reassortant virions from el Hussein et al. and Samal et al. [REFs]*

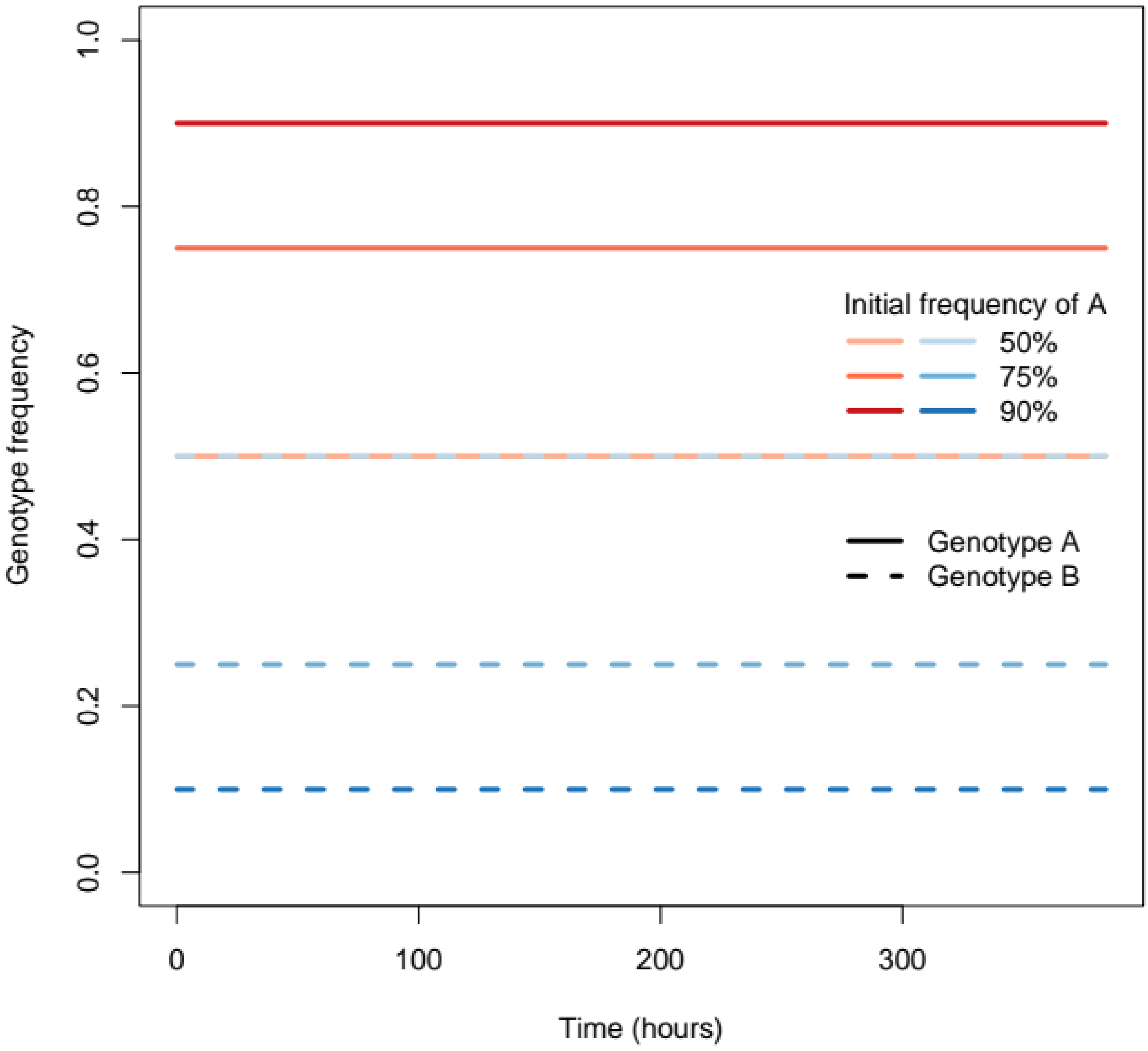


*Fig. S2. If both genotypes have the same parameters, then each exhibits stable frequencies under coinfection without reassortment. I.e., there is no a priori stable strain frequency built into the model.*

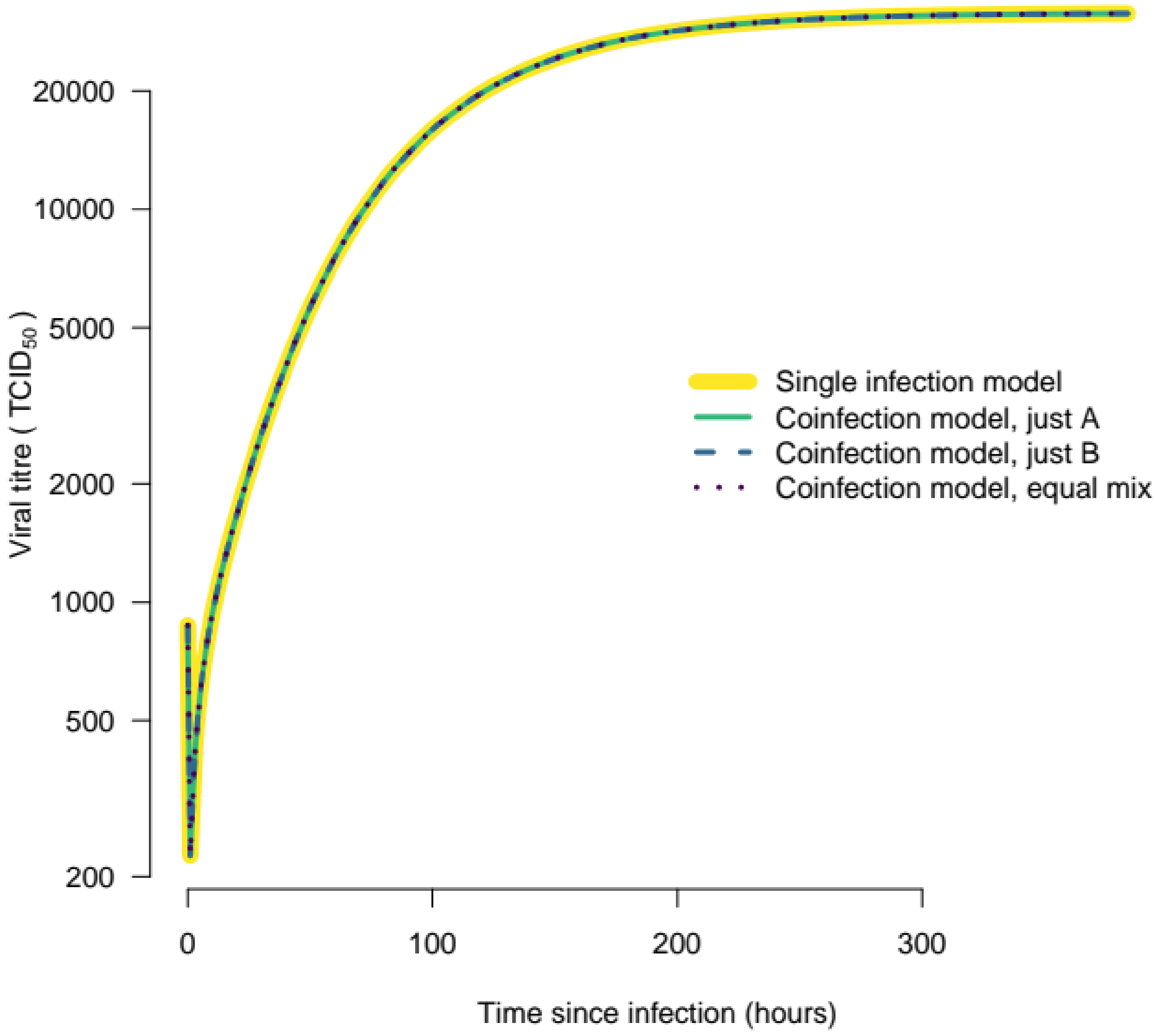


*Fig. S3: In the absence of reassortment, the single model and the coinfection model behave identically, irrespective of the initial mix of virus types in the bloodmeal (and assuming both types have the same parameters)*

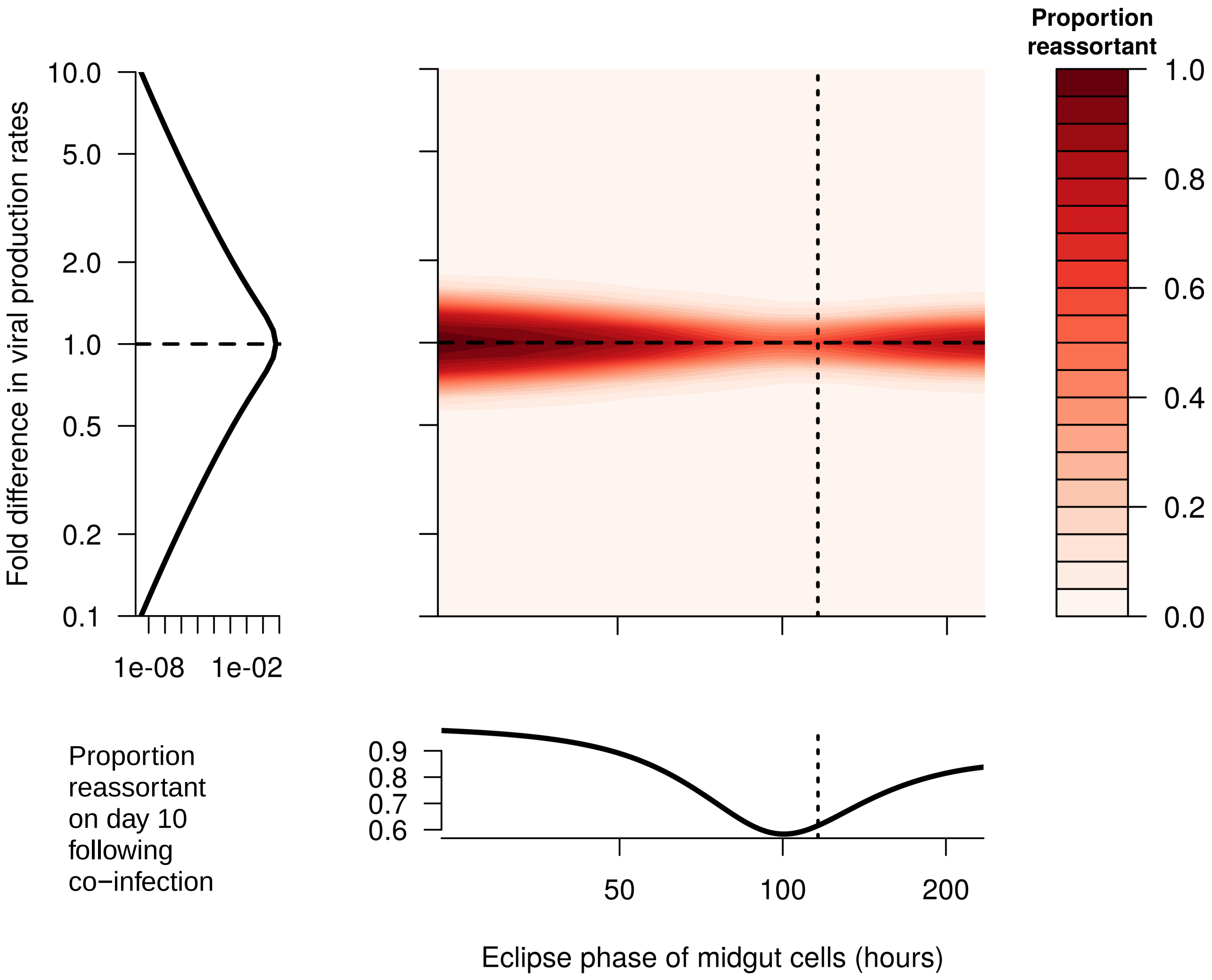


*Fig. S4: A: The proportion of reassortant viruses observed ten days after the initial infection (color-scale) when the blood meals occur on day 0, for different lengths of eclipse phase (x-axis) and for different viral production rates between the two infections (y-axis). When the fold-change in p is less than 1, the first infecting virus has higher production rate than the second, and when it is greater 1 the second virus has higher production rate. Dashed lines represent baseline values. Note that the eclipse phase in the secondary tissues was also varied by proportionally the same amount from baseline as the eclipse phase in the midgut (not shown).*

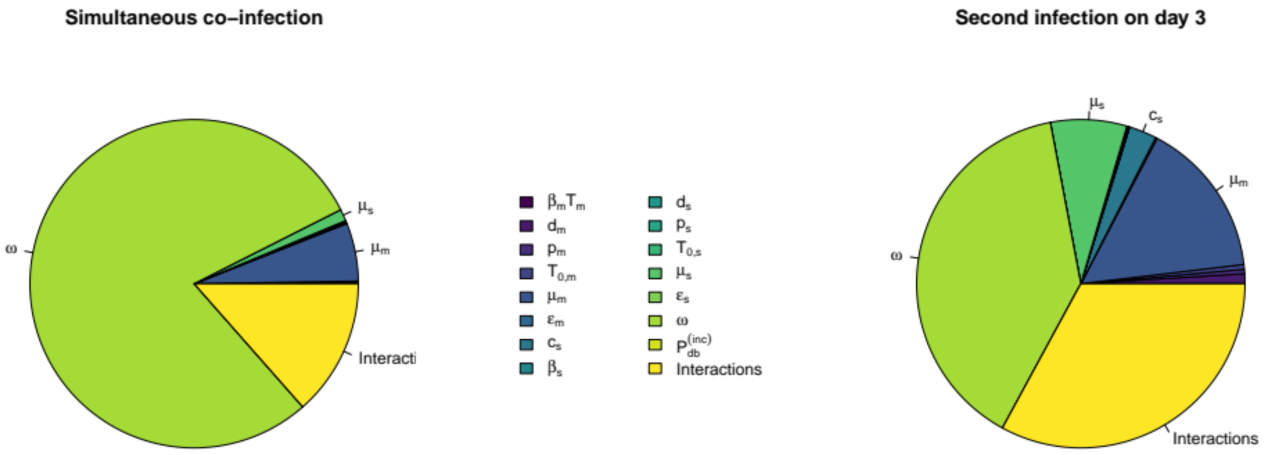


*Fig. S5: First-order indices for Saltelli variance-based sensitivity analysis of the proportion of viruses that are reassortant. Pie segments represent the variance explained by each parameter on its own, with the final segment describing the amount of variance explained by interactions between parameters. The most important parameters are labelled: ω is the proportion of virions produced by a coinfected cell that are with-like (i.e. are not reassortants), μ is the rate of cellular death, and c is the rate of viral clearance/death. Left panel shows the indices for the proportion reassortant on day 10 following a simultaneous coinfection on day 0. Right panel is the same, but for infection on day 3.*

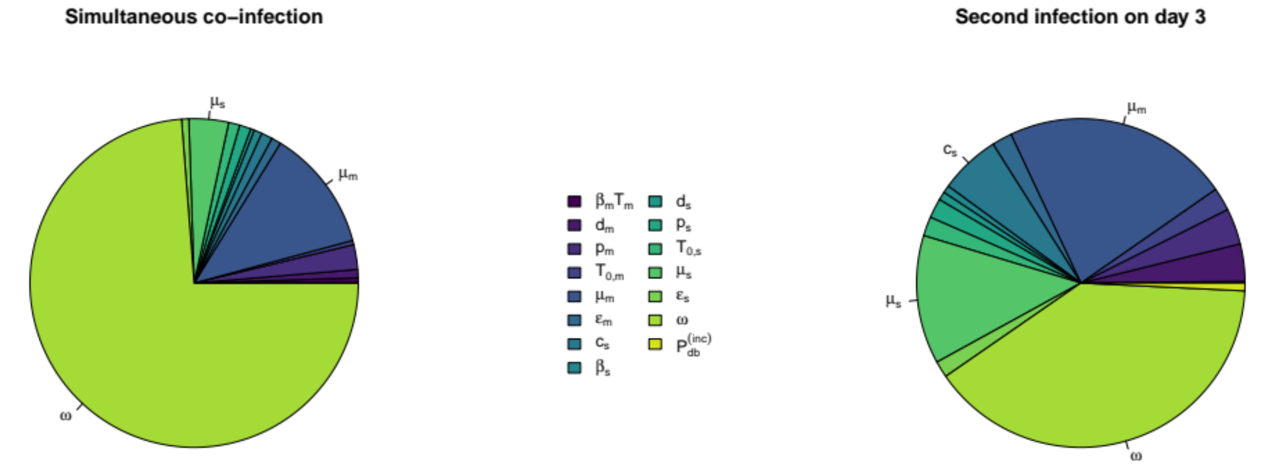


*Fig. S6: Total-order indices for Saltelli variance-based sensitivity analysis of the proportion of viruses that are reassortant. Pie segments represent the variance explained by each parameter, both on its own*

*and in interactions with all other parameters. The most important parameters are labelled: ω is the proportion of virions produced by a coinfected cell that are with-like (i.e. are not reassortants), μ is the rate of cellular death, and c is the rate of viral clearance/death. Left panel shows the indices for the proportion reassortant on day 10 following a simultaneous coinfection on day 0. Right panel is the same, but for infection on day 3.*
